# Electrical stimulation elicits space- and parameter-dependent spiking responses in human cortical organoids

**DOI:** 10.1101/2025.09.01.673532

**Authors:** N. Seseri, D. Nigrisoli, F. Faraci, A. D’Angelo, R. Freddi, S. Chandran, R. Barbieri, S. Corti, L. Ottoboni, S. Russo

## Abstract

Electrical stimulation (ES) is used to treat neuropsychiatric disorders and investigate brain dynamics, yet its effects on human cortical microcircuits remain poorly understood. Cortical organoids provide a unique platform to investigate these mechanisms in isolation from subcortical and long-range cortical inputs. Here we illustrate how cortical organoids respond to ES, identifying the response profiles of isolated cortical circuits while detailing a roadmap of how ES parameters affect the organoid spiking activity.

We employed a high-density multielectrode array to record neuronal activity from cortical organoids (n=417 units in N=7 organoids) during ES, systematically varying stimulation frequency, intensity, pulse width, and charge density.

By analyzing single unit spiking activity, we found that ES elicits excitatory, inhibitory, and mixed responses in 39%, 12%, and 17% of the units, respectively. On average, this response lasted 100 ms and became stable within 26 trials. The magnitude of both excitatory and inhibitory responses was maximal near the stimulation site and decayed with distance. The response magnitude was inversely correlated with pulse intensity and duration, but not with stimulation frequency and charge density.

These findings demonstrate that local cortical circuits are sufficient to initiate the early excitatory phase of the canonical ES response, whose magnitude depends on ES parameters, and can sustain the excitatory phase for over 100 ms. The reduced late inhibitory phase, together with the absence of late excitatory components observed 200 ms after ES in intact adult brains in-vivo, suggests that these phases may depend on neuronal maturation or inter-area connections. Our work thus establishes cortical organoids as a framework for studying the local contributions to ES-induced activity in a developmental model of the human cortex.

## INTRODUCTION

Cortical stimulation enables testing and modulating physiological and pathological brain dynamics^1–3^, yet the specific underlying neuronal circuits it activates remain unclear ^4^. At mesoscopic and macroscopic scales, combining electrical stimulation (ES) with electrophysiological recordings has revealed how a local stimulus influences the activity of connected areas^5,6^. Over the past decades, this approach has been extensively used to quantify state-dependent changes in human brain dynamics across wakefulness and sleep^7,8^, sensorimotor tasks^9^, brain injuries^10^, and epilepsy^11,12^. However, ES has often been delivered according to empirical or clinical criteria, thus leading to substantial heterogeneity in its parameters and in the resulting effects.

Recent systematic studies have contributed to clarifying how specific ES parameters—such as intensity, pulse duration, and frequency—shape neural responses in humans^13–16^. These findings reveal that ES parameters directly impact the macroscopic and mesoscopic effects of ES, which suggests that these changes may originate from a different engagement of cortical microcircuits.

Although technological advancements have enabled direct recordings from single neuron responses to ES in humans^2,17–21^, the invasive nature of these methods restricts their use to patients with neurological diseases and potentially pathological circuits. An alternative approach is to infer the microcircuits engaged by ES using multispecies experimental designs ^9^, although this indirect approach requires careful control for potential confounds across species. A more direct approach is represented by testing ES effects in human brain slice cultures obtained from spare CNS tissue from neurosurgical procedures^22,23^,24. However, this approach is restricted to tissue derived from clinical or post-mortem cases that substantially constrain its translatability to in-vivo models. While the effects of ES can be also studied on traditional neuronal cultures derived from human induced pluripotent stem cells (hiPSC)^25^, these models exhibit substantial differences with the three dimensional architecture of the human brain^26^.

These limitations could be overcome by cortical organoids obtained from hiPSC^27,28^, offering a model that not only recapitulates human cortical architecture and development, but also enables long-term monitoring and controlled causal manipulations^29–33^. Yet, using organoids to study brain dynamics requires testing whether, and to what extent, their electrical activity resembles the one of physiological brains^34,35^.

Neural organoids have been shown to replicate key phenotypic characteristics of the human brain, including omic and biochemical signatures of specific brain areas^36–38^ as well as connectivity capable of generating population-level oscillations^39^. This cytologic and architectural substrate makes them a promising model for studying human electrophysiological activity under controlled conditions^27,40^.

Previous research found key similarities between the electrophysiological properties of cortical organoids and the human brain. Specifically, the spontaneous electrical activity of cortical organoids evolves over maturation time, transitioning from periodic bursts of activity at ∼2 months, to more rapid nested oscillations at 6-8 months, reminiscent of the electroencephalographic (EEG) pattern of preterm newborns^39,41^. Furthermore, their NMDA and GABA receptor maturation parallels the one observed in the postnatal development of the human cortex^42,43^. The similarities between organoid activity and known electrophysiological profiles of brain development suggest that they may serve as a viable model system for studying human cortical dynamics throughout brain development^44^.

Furthermore, neural organoids are currently being explored as a bio-computation platform with the aim of developing new highly efficient computational architectures^27,45^. These studies combine high-density microelectrode arrays (hd-MEA) and ES to mediate closed loop communication between neural organoids and artificial neural networks^46^. This combination has enabled unsupervised learning in neural organoids, inducing short-term plasticity, reconfiguring the organoid functional connectivity, and showcasing the potential for developing complex organoid-computer interfaces^27^. Beyond computation, specific ES patterns (e.g. theta-burst stimulation) proved effective in affecting early gene expression, criticality measures, neuronal network dynamics, and synaptic plasticity^47^. Notably, these effects can be pharmacologically modulated to mirror synaptic modulation and short-term potentiation, further consolidating organoids as a platform for studying human neurophysiological processes. Despite the remarkable potential of organoids ES applied to drug development, neurophysiology research, and biocomputing, to date the effects of ES -and its parameters-on neural organoids remains relatively unexplored.

In this study, we characterize the electrophysiological response of cortical organoids to ES across spatial and temporal domains. We recorded the neuronal spiking activity evoked by a series of electrical pulses while systematically varying pulse amplitude and duration, stimulation frequency, and charge density. We found that electrical stimulation induced excitatory, inhibitory, and mixed effects that are maximally prominent nearby the stimulation site. Population-wise, ES elicited a prominent increase in firing rate, whose magnitude was inversely correlated with ES pulse amplitude and duration. We provide a systematic characterization of the temporal, spatial, and functional properties of the electrophysiological response of cortical organoids to ES. Compared to the typical triphasic excitation-inhibition-excitation pattern evoked by ES in vivo, our results reveal that ES-induced early excitation is preserved in cortical organoids, while the intermediate inhibition is reduced and the late rebound excitation is lost. These results draw a roadmap to link distinct phases of the ES response to their neurophysiological substrate and to guide the selection of ES parameters in cortical organoids.

## METHODS

### Ethical approval

This study was conducted in accordance with the ethical principles of the Declaration of Helsinki and with institutional and national regulatory guidelines. Written informed consent was obtained from all participants, as approved by the local ethics committees, for the collection, storage, and analysis of biological samples and clinical data. All experiments complied with international GLP and GCP standards and regulations.

### Generation and maintenance of iPSC lines

For iPSC line #1, peripheral blood mononuclear cells (PBMCs) were isolated from a healthy control subject using Ficoll-Paque density gradient centrifugation (age=64, sex=F). PBMCs were reprogrammed using the Sendai virus delivery system (CytoTune™ iPS 2.0, Thermo Fisher Scientific) according to the manufacturer’s instructions^48^. iPSC colonies were selected based on morphology and expanded on Cultrex-coated plates (R&D Systems) in mTeSR1 medium (Stem Cell Technologies). Cell cultures were maintained at 37°C, 5% CO2, with daily medium change. iPSC lines were passed using EDTA (0.5 uM) for dissociation. iPSCs were controlled for chromosomal abnormalities by karyotyping, pluripotency assessed by immunofluorescence staining for OCT4, SOX2, NANOG, and TRA-1-60, and by quantitative RT-PCR analysis of pluripotency markers^49^.

iPSC line #2, was an isogenic line generated in Siddharthan Chandran’s Laboratory under full Ethical and Institutional Review Board approval of the University of Edinburgh^50^.

### Generation of Cortical Organoids

hCOs were generated following the protocol described by Miura et al.^51^. Briefly, iPSC colonies were dissociated into single cells using Accutase (Thermo Fisher Scientific) and seeded at a density of 10,000 cells/well in ultra-low attachment 96-well plates with mTeSR1 medium (StemCell Technologies) supplemented with Y27632 ROCK inhibitor (10 μM, Cell Signaling) for 24 hours to promote embryoid body (EB) formation. From day 0 to day 6, the medium was supplemented with dorsomorphin (DM, 5 µM, Sigma-Aldrich), SB-431542 (10 µM), and XAV-939 (1.25 µM, Tocris). At day 6, cultures were switched to neural differentiation medium (Neurobasal medium; 2% B27 supplement without vitamin A, Thermo Fisher Scientific; 1% Glutamax; 1% penicillin-streptomycin) supplemented with epidermal growth factor (EGF, 20 ng/mL, Peprotech) and basic fibroblast growth factor (bFGF, 20 ng/mL, Peprotech). On day 12, EBs were transferred to low-attachment 24-well plates, with medium changes every two days. From day 22 onward, EGF and bFGF were replaced with BDNF (20 ng/mL), neurotrophin-3 (NT3, 20 ng/mL, Peprotech), ascorbic acid (0.2 mM), dibutyryl cyclic AMP (d-cAMP, 50 µM, Sigma-Merck), and docosahexaenoic acid (DHA, 10 µM, Merck). From day 46 until the end of differentiation, hCOs were maintained in neural maintenance medium (Neurobasal medium; 2% B27 Plus supplement, Thermo Fisher Scientific; 1% Glutamax; 1% penicillin-streptomycin). Terminal differentiation was carried out for up to 310 ± 67 days, with medium changes every 4–5 days. Organoid size and morphology were monitored weekly, and those showing necrosis or abnormal development were excluded from analysis.

### Immunofluorescence

Organoids needed for characterization were fixed in cold 4% paraformaldehyde for 20 minutes, embedded in a 7.5% sucrose/10% gelatin (Gelatin from bovin, Merck) solution for cryopreservation and then cryosectioned at 14 μm thickness. Cut slices were preserved at - 80 °C until staining. Blocking with a solution containing 5% normal goat/donkey serum, 5% BSA, and 0.3% Triton X-100 for 1 hour at RT was performed, primary antibodies were applied overnight at 4°C for characterization. Secondary antibodies included Alexa Fluor 488-, 568-, and 647-conjugated goat anti-mouse, anti-rabbit, and anti-rat antibodies (Thermo Fisher Scientific) used at 1:1000 dilution and were applied for 1 hour and a half at RT. Images were acquired using a Leica SP8 confocal microscope.

### Electrophysiological recording protocol

Between 2 and 7 days before the recording, organoids were moved to BrainPhys™ Neuronal Medium with SM1 Kit (Stemcell Technologies) integrated with small molecules as per the differentiation protocol. Accura Chip (3Brain AG), the day of the experiment, was washed and gently brushed with PBS to remove air bubbles and reach consistent detection on the chip. The chip noise background was tested recording on a Biocam Duplex (3Brain AG) using BrainWave5.0 software, and the chip excluded when the noise level exceeded +/-50 μV, which would indicate hydrophobicity. In case of high noise level exceeding this threshold, PBS was removed, and we repeated washes with a brush and 70% Ethanol, the chip was rinsed in water, and PBS was reintroduced. Afterwards, PBS was removed, the medium was placed on the chip with the organoid. Excess medium was removed with a 10μl pipette. The organoid was secured to the surface of the electrodes using a plastic anchor and the chamber was filled with medium. The chip was then inserted into the BioCam Duplex and the platform heating was activated with a target temperature of 37°C.

### Electrophysiologic data acquisition

Data were acquired using a 4096 channels multielectrode array (BioCAM DupleX system, 3Brain AG) with a sampling rate of 19754 Hz and a hardware high pass filter at 100Hz (electrode size: 21×21 μm, pitch: 60μm). Raw data was recorded using BrainWave5.0. We acquired spontaneous activity and, before starting the electrical stimulation protocol, 4-aminopyrdine (4AP; 50umM) was added to the recording chamber.

### Electrical stimulation

To maximize the probability of effectively engaging neurons, the stimulation site was visually selected by the operator as two close groups of 3 MEA channels (each group separated by one line of channels) located in the area of maximum spiking activity. The stimulation was administered as a bipolar, biphasic stimulation, by passing current between the two groups of channels, one positive and one negative. We set the default stimulation parameters to pulses of 90 μA intensity and 200 μs duration delivered across 6 channels (3 at each pole) at a frequency of 1 Hz. Each stimulation parameter was systematically changed while keeping all the other parameters fixed. For example, the effect of stimulation intensity was investigated by administering 5 trains (i.e. 30, 60, 90, 120, 150 μA) of 30 stimuli each, with an inter-train-interval of 1-2 minutes. An analogous protocol was applied for pulse duration (50, 100, 200, and 300 μs), frequency (0.2, 0.5, 1, and 3 Hz), and charge density (2, 3, and 4 channels per pole). This protocol led to 30 stimuli per each variation of the stimulation parameters, and only for the stimulation with default parameters that was repeated 4 times a total of 120 stimuli. For each experiment, the order of the stimulation parameters was defined as follows: first, we randomized the order of the explored parameters, and for each parameter, we randomized the order of the applied values. This randomization maximized comparability within variations of the same parameter while preventing biased effects due to the order of stimuli. Additionally, it provided several repetitions of the default parameter set to assess whether the responses to the same stimulation changed over time.

For each organoid, at the end of all the recording sessions, we administered tetrodotoxin (TTX) and verified that spiking activity and responses were abolished.

### Data preprocessing

BrainWave data were imported and preprocessed using SpikeInterface and custom python scripts. The artifacts associated with the electric stimulation were removed with a convolution-based method compatible with the SpikeInterface framework^52^. The algorithm operates in two sequential stages. First, artifacts were identified by convolving each channel’s signal with a pre-defined artifact template and identifying signals that, in a given time window exceeded 5 std of the overall trace and the absolute threshold of 1×10^6^ μV^2. Then, artifacts were removed by replacing the identified artifact segments (duration 7 ± 3 ms) with linear interpolation between boundary values, with added Gaussian noise (σ = 20μV) to preserve signal statistics. The recording was then bandpass filtered from 300 to 6000 Hz.

Units were sorted using Kilosort 2.5 ^53^. Sorted units were prefiltered by selecting those with an inter spike interval (ISI) violation ratio below 10% (percentage of ISIs < 2 ms) and excluding units in which more than 30% of inter-spike intervals (ISIs) clustered around multiples of 0.4 s, a pattern consistent with MEA calibration artifacts. The remaining units were manually curated using Phy.

### Data preprocessing and processing

For each unit and set of stimulation parameters, we computed the peristimulus time histogram (PSTH) from −300 ms to +500 ms relative to the time of ES.

Among all the identified units, we considered for subsequent analyses only those that exhibited at least one spike (either before or after the stimulus) in 30% of the 120 ES trials with default ES parameters, that we named active units. The PSTH was smoothed with a Gaussian filter (σ = 7 ms) to reduce noise before analysis. For visualization purposes only, the blanked segment of the signal (±5 ms around stimulus onset) was replaced with a mirrored copy of the signal from −15 ms to −5 ms before stimulus onset.

To characterize the heterogeneity of excitatory and inhibitory patterns across units over time, we applied the criteria for identifying excited and inhibited units with time bins of 25 ms. Excitation and inhibition thresholds were computed based on pre-stimulus activity: the excitation threshold was defined as the mean plus 5 std of the baseline distribution, while the inhibition threshold was defined as the 5th percentile value of the baseline distribution. Units were classified as excited if their smoothed response exceeded the excitation threshold, and as inhibited if their response dropped below the inhibition threshold for two consecutive time bins. Units showing both features at different time bins were classified as mixed. Mixed units were further classified based on the first modulation identified as “excited-inhibited” or “inhibited-excited”.

To determine the minimum number of trials required to obtain reliable PSTHs, we used ES with default parameters to compare PSTHs computed with varying trial counts (1-120) against the reference PSTH derived from all 120 trials. Reliability was assessed using cosine similarity, mean squared error (MSE), and Pearson Correlation. For each trial count, each measure was averaged across units. The minimum number of trials required to obtain a reliable PSTH was defined as the smallest trail count yielding a cosine similarity ≥0.9 between the PSTH computed on n trials and the 120-trial reference PSTH. To ensure robustness, each measure was repeated 100 times per trial count, with trials randomly selected in each iteration. For visualization purposes, MSE was normalized as a percentage of the average MSE at trial 1.

We also assessed how the variability across neurons changed over time after ES. To this end, we computed the Fano factor, defined as the variance to mean ratio of the spike counts ^54^. For each neuron, spike trains were aligned to stimulus onset and spike counts were obtained using a 10-ms sliding window with a 5-ms step. To avoid contamination by stimulus-related artifacts, signals 5 ms before or after ES time were excluded. For each window, we calculated the Fano factor across trials and then averaged the resulting time courses across all responsive neurons. Finally, we compared the post-stimulus trajectory of the Fano factor computed on all the neuronal population to its pre-stimulus baseline level and defined the recovery time as the first point after stimulus onset when the mean Fano factor returned within the 95% confidence interval of the baseline.

To determine the onset, offset, peak, and switch latency of the PSTH, we analyzed the smoothed PSTH (Gaussian kernel with σ = 7 ms). The peak latency was identified as the time of the maximum response within 5 and 500 ms after the stimulus onset. Onset and offset times were defined as the PSTH crossings with the mean (or 5th percentile) of the baseline (from −300 to −5 ms) closest to the peak (or through). Switch time was defined as the difference between the offset of the first event and onset of the second event.

Then, we assessed the impact of distinct stimulation parameters on the organoid spiking response. To compare across parameter values accounting for variability of the baseline activity and responsiveness across neurons, we normalized the response of each set of parameters to the default set of parameters. To provide an exhaustive picture of the effects of each parameter, this was done both by subtracting and computing the percentage variation compared to the spiking response elicited by the default parameter set.

Data were analyzed using Friedman test for repeated measures and Wilcoxon tests for post hoc analysis, with multiple comparison correction using the False Discovery Rate method from Benjamini and Hochberg.

## RESULTS

We recorded the spiking response evoked by the electrical stimulation of brain organoids aged 310 ± 67 days. In 7 organoids, we isolated 417 single units (59 ± 40 per organoid), of which 197 were active units (i.e. 1 or more spikes in 30% of the trials; Figure 1A-B) and 135 (68.6%) of the latter exhibited a modulation of the firing rate in response to ES (90μA, 1Hz, 200μs, 6 channels) within 500 ms from the stimulus onset (Figure 1 C-D).

**Figure 1.**
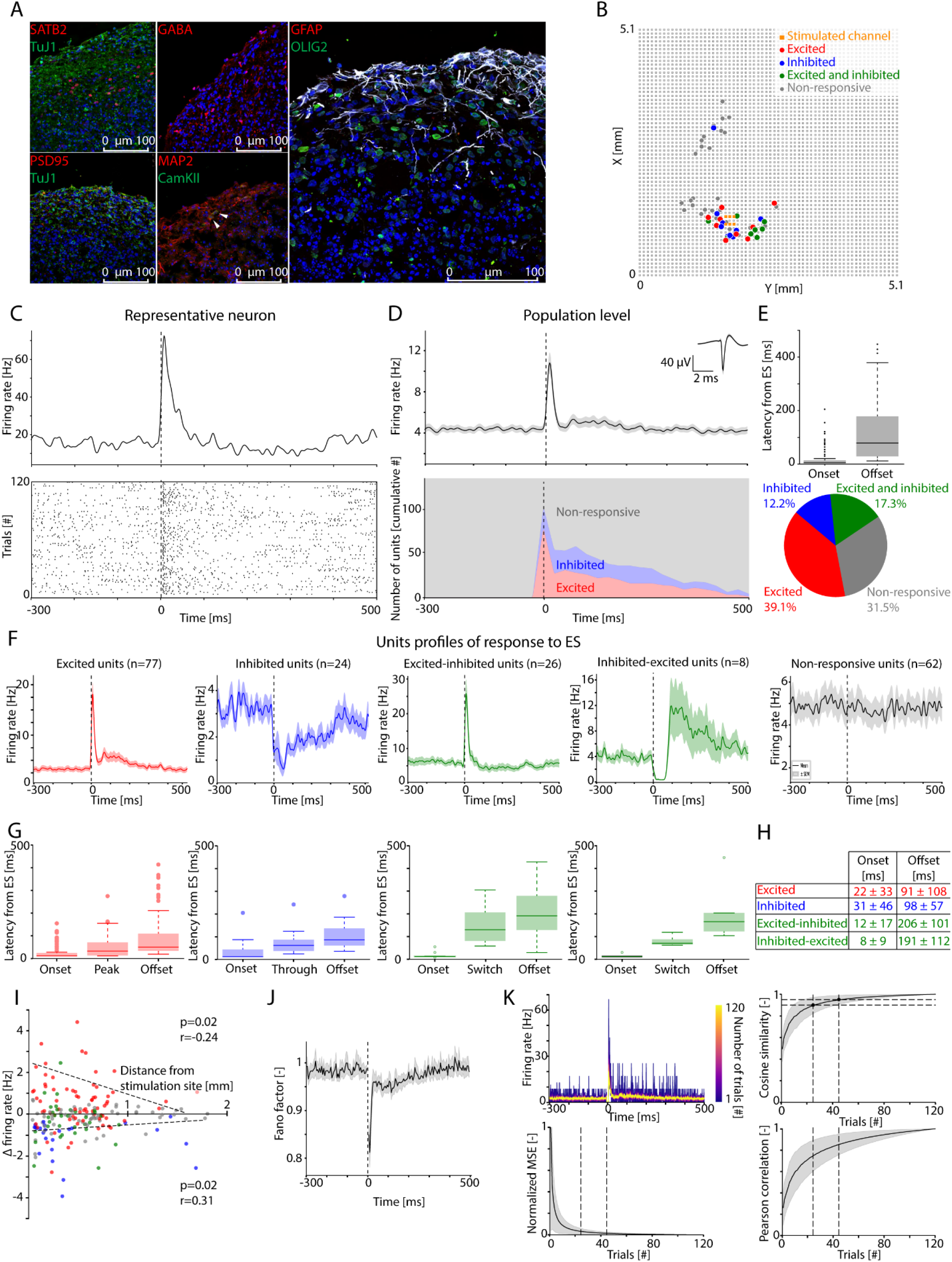
Cortical organoids respond to electrical stimulation with excited and inhibited spiking patterns. **Panel A -** Representative Immunofluorescence images of human Cortical Organoids at day 125 of differentiation showing the expression of neural markers (Tuj1, MAP2), upper-layer cortical markers (SATB2), glial markers (GFAP, OLIG2), excitatory marker (CamKII, white arrow), inhibitory marker (GABA) and post-synaptic marker (PSD95). Nuclei were stained with DAPI (blue). Scale bar: 100 μm. **Panel B -** Reconstruction of the hd-MEA electrodes showing, for a representative organoid, the location on the stimulating electrodes (orange squares) with respect to the recorded units (Color coded based on ES response as in Panel E). **Panel C -** Peri-stimulus time histogram (PSTH, top) and raster plot (bottom) of the response to ES from a representative neuron. **Panel D -** Top panel: Average PSTH of the response to ES across all responsive units. Bottom panel: Time course of the number of the units excited, inhibited, or non-responsive to ES for each 25 ms time bin. **Panel E -** Top: Boxplot showing the time of ES-induced response onset and offset across all responsive units. Bottom: Pie chart showing the percentage of units in which ES induced excitation (red), inhibition (blue), both excitation and inhibition (green), and no response (gray). **Panel F -** Average PSTH across each group of units: excited (red), inhibited (blue), first excited then inhibited (green left), first inhibited then excited (green right), and non-responsive (gray). **Panel G -** Boxplot showing, for each group of units, the time of onset and offset of the PSTH modulation, together with the time of maximum modulation (peak and through for excited and inhibited units, respectively), and the time of switch between excited and inhibited phases (for excited-inhibited and inhibited-excited units). **Panel I -** Scatter plot and linear fit showing the relationship between the change in firing rate compared to baseline and the distance from the ES site (color code as in Panel E; 6 outliers were removed for visualization purposes). The graph shows a negative relationship between the magnitude of the firing rate modulation and the distance from the ES site for both positive (r= −0.24; p=0.02) and negative modulations (r=0.31; p=0.02). **Panel J -** Time course of the Fano Factor (FF) over time showing a short-lived, large FF reduction immediately after the stimulus onset, followed by a more subtle and longer-lasting FF reduction. **Panel K -** Top left panel: Trace showing the PSTH computed across neurons from increasing numbers of trials. Top right panel: Plot showing cosine similarity (average - black trace; standard deviation: shaded gray area) as a function of the number of trials. The dashed lines indicate the 0.9 and 0.95 thresholds of the cosine similarity (horizontal lines) and their corresponding number of trials (vertical lines). Bottom left panel: Plot showing how the Mean Square Error (MSE; average: black trace; standard deviation: shaded gray area) changes as a function of the number of trials. The dashed vertical lines indicate the number of trials identified from 0.9 and 0.95 cosine similarity. Bottom right panel: Plot showing how Spearman correlation of the response computed from 120 trials (average: black trace; standard deviation: shaded gray area) changes as a function of the number of trials. The dashed vertical lines indicate the number of trials identified from 0.9 and 0.95 cosine similarity.

This modulation assessed throughout the response window reflects the sum of heterogenous effects that change over time and across units. Time-wise, ES immediately elicited transient effects: both the number of excited and inhibited units was maximal immediately after the stimulus onset and progressively returned to baseline levels in ∼100 ms, with some units being modulated for a longer time (Figure 1D-E). Unit-wise, this quantification aggregates excitatory and inhibitory modulations across units: 39.1% of the units were excited, 12.2% were inhibited, 17.3% were both inhibited and excited, and 31.5% did not respond to ES (Figure 1E).

This response pattern at population-level reflects heterogeneous single unit responses. To interpret these heterogeneous responses, we clustered units based on their response profile: excited, inhibited, first excited then inhibited (excited-inhibited), first inhibited then excited (inhibited-excited), and non-responsive. For each cluster we computed its PSTH (Figure 1F), finding that both excited and excited-inhibited units exhibit a large transient increase in spiking rate followed by a second shallow excitation in the former and by a shallow inhibition in the latter. By contrast, inhibited units exhibited an abrupt reduction of firing rate that progressively returned to baseline levels, while inhibited-excited units showed an initial deep inhibition followed by a prolonged excitatory response.

Quantitatively, all response onsets occurred almost immediately after ES (20 ± 33 ms) while offsets were more variable, with the earliest offset observed in excited units 12 ms after ES, and the latest offset observed in excited-inhibited units after 448 ms (see Figure 1G-H).

We wondered whether this heterogeneity across neuronal spiking responses was influenced by biophysical parameters, such as neuronal spatial distribution. To answer this question, we explored whether excitatory and inhibitory spiking modulations were explained by their distance from the ES site. We found that the relative magnitude of both excitatory (Spearman correlation; r=-0.24; p=0.0187) and inhibitory (Spearman correlation; r=0.31; p=0.00169) responses were correlated with the distance from the ES site and fell to baseline levels within 2 mm from the ES site (Figure 1I).

Given the heterogeneity in spiking responses to ES, we assessed whether ES could enhance variability also in the spiking activity across neurons, as measured by variability with respect to a hypothetical Poisson distribution. Thus, we investigated how ES affected Fano Factor across neurons, finding that ES induced a large, immediate and short-lived drop of the Fano factor, followed by a longer reduction of Fano factor lasting 280 ms (Figure 1J).

This heterogeneity across neuronal responses prompted us to investigate the extent of trial-to-trial variability in ES responses and to determine how many trials were required to obtain a reliable response. To this end, we evaluated how the averages of progressively higher numbers of trials approximate a ground truth response, obtained from averaging 120 trials (Figure 1K). We found that 26 trials were sufficient to achieve a cosine similarity value of 0.9, corresponding to a Pearson correlation and a mean square error (MSE) compared to the ground truth response of 0.76 and 0.03 (∼3%), respectively. 47 trials were required to achieve a cosine similarity value of 0.95, corresponding to a Pearson correlation and an MSE compared to the ground truth response of 0.86 and 0.01 (∼1%), respectively. To reach 0.9 and 0.95 of the Spearman correlation required 60 and 84 trials, respectively. To achieve 10%, 5%, and 1% of the MSE required 9, 17, and 54 trials, respectively.

Since ES parameters can significantly affect the properties of the neural response ^15,16^, we explored whether -and how-distinct ES parameters independently shaped the evoked spiking response. To this end, we independently explored the effect of pulse intensity and duration, stimulation frequency, and charge density on evoked spiking while keeping the other parameters fixed. To compare the effect of stimulation parameters while accounting for variability across neurons, we computed the effect of each parameter on spiking activity relative to the spiking activity elicited by a fixed set of stimulation parameters (90μA, 1Hz, 200μs, 6 channels; as in Figure 1).

To assess the effect of pulse intensity, each stimulus was administered at 30, 60, 90, 120, and 150 μA. We found that lower stimulation intensities were associated with enhanced spiking responses, while higher stimulation intensities were associated with reduced spiking responses (Friedman Test; p<0.001; Figure 2A). Compared to 90μA (5.96 ± 6.98 Hz), 30 μA pulses induced 31.92% more spikes (7.87 ± 8.24 Hz), 60 μA pulses induced 20.71% more spikes (7.20 ± 7.38 Hz), 120 μA pulses induced 0.54% more spikes (6.00 ± 6.95 Hz), and 150 μA pulses induced 10.48% less spikes (5.34 ± 7.09 Hz). This reflects a linear decreasing relationship between stimulation intensity and spike number, with a decrease of 0.02 Hz/μA (0.27% per μA).

**Figure 2.**
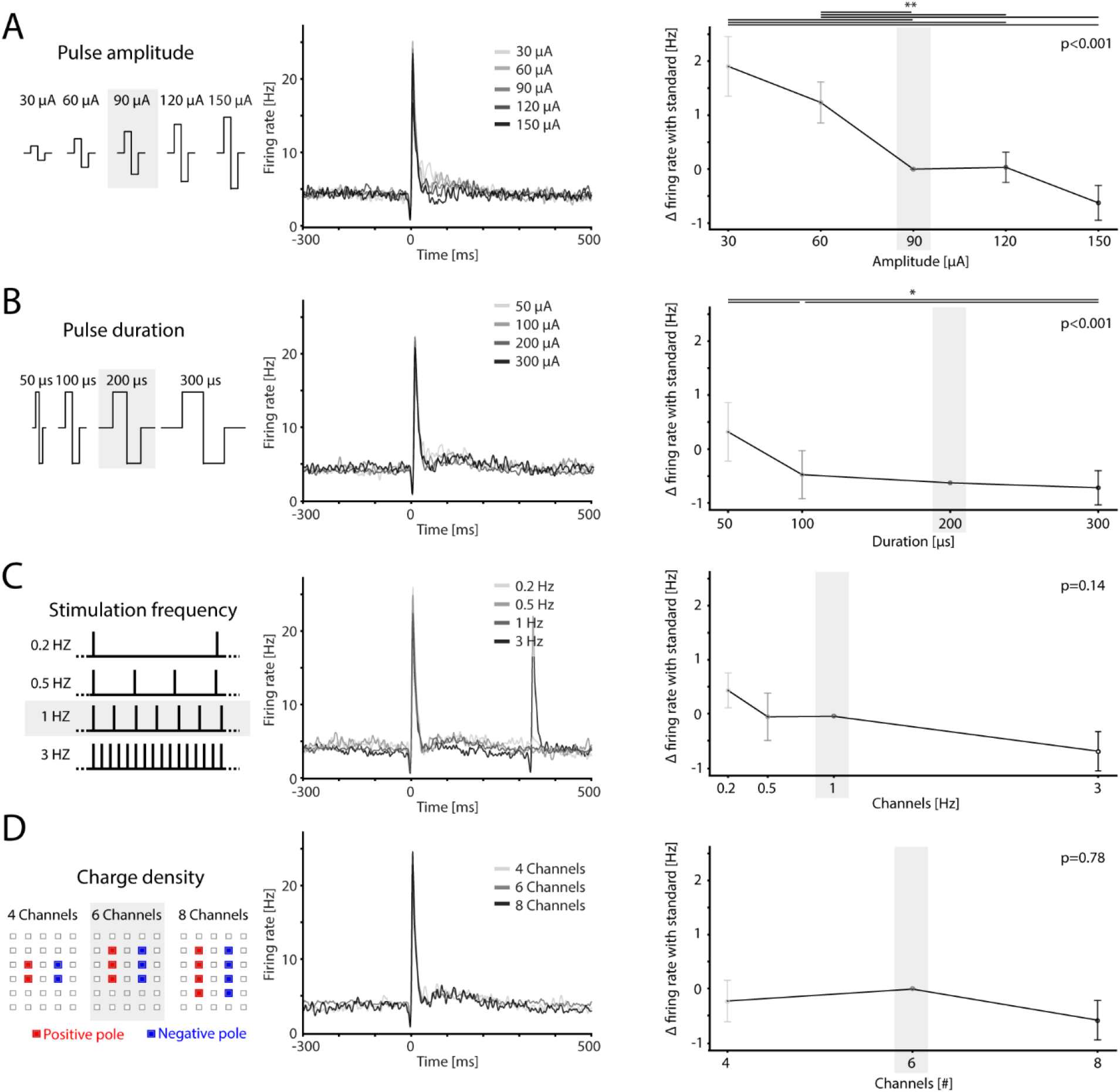
Cortical organoid responses to ES depend on ES biophysical parameters. **Panel A -** Left to right: schematic representation of the different pulse amplitudes. PSTH of a representative neuron for each pulse amplitude (gray shades). Average PSTH across all neurons for each pulse amplitude. The error plot shows mean (circles) and standard error (bars) of the difference of firing rate between the ES response to each pulse amplitude and to the default parameters (Friedman Test; p<0.001; post-hoc comparisons are reported in figure: *: p≤0.05; **: p≤0.01; no line: p>0.05). The default set of parameters is highlighted by the gray shaded area. **Panel B -** Same as Panel A for different values of pulse duration (Friedman Test; p<0.001). **Panel C -** Same as Panel A for different values of stimulation frequency (Friedman Test; p=0.14). **Panel D -** Same as Panel A for different charge densities, achieved through different numbers of stimulating channels (Friedman Test; p=0.78).

To assess the effect of pulse duration, each stimulus was administered with a width of 50, 100, 200, and 300 μs. We found that lower pulse duration values were associated with larger spiking responses (Friedman Test; p<0.001; Figure 2B). Compared to 200 μs (6.91 ± 7.04 Hz), 50 μs pulses induced 13.48% more spikes (7.84 ± 8.40 Hz), 100 μs pulses induced 2.16% more spikes (7.06 ± 7.14 Hz), 300 μs pulses induced 1.30% less spikes (6.82 ± 8.05 Hz).

To assess the effect of stimulation frequency, each stimulus series was administered with a frequency of 0.2, 0.5, 1, and 3 Hz. We did not observe a significant effect of stimulation frequency on spiking rate (Friedman Test; p=0.14; Figure 2C), even though there was a trend suggesting that higher stimulation frequencies may be associated with reduced spiking responses. Namely, compared to 1 Hz stimulation frequency whose average response frequency was 6.51 ± 7.40 Hz, there was a 7.35% increase with 0.2 Hz (6.99 ± 7.32 Hz), 0.13% reduction with 0.5 Hz (6.50 ± 7.33 Hz), 9.88% reduction with 3 Hz (5.87 ± 8.01 Hz).

To assess the effect of charge density, each stimulus was administered either between 4, 6, or 8 channels. We did not observe a significant effect of the number of stimulating channels on the evoked spiking rate (Friedman Test; p=0.78; Figure 2D). In detail, compared to the 6-channel stimulation (6.86 ± 7.43 Hz), there was a 3.18% decrease when using 4 channels (6.65 ± 7.83 Hz) and 8.28% decrease when using 8 channels (6.30 ± 7.38 Hz).

Besides minimizing the effect of session order and time on neural activity, the hierarchical randomization across stimulation series enabled us to evaluate the impact of time over ES responses by comparing the responses evoked by identical stimulation parameters at the beginning and end of the experiment. The first and last series of the standard ES, delivered at 14.13 ± 1.76 minutes of distance, evoked spiking responses that were not statistically different (first ES series: 4.62 ± 4.63 Hz; last ES: 4.60 ± 4.93 Hz; Wilcoxon Signed Rank Test; p=0.81).

## DISCUSSION

Our study shows that ES of cortical organoids elicits a mixed but mainly excitatory modulation of their neuronal firing rate that, on average, starts 21 ms after the stimulus onset and lasts 101 ms. The magnitude of this excitatory response negatively correlated with the intensity and duration of the electrical stimulus but did not correlate with its frequency and charge density. The magnitude of both excitatory and inhibitory effects of ES were negatively correlated with the distance from the ES site. Furthermore, 26 trials were sufficient to compute reliable average responses, with a 90% similarity with the response computed over 120 trials.

Organoids responded to electrical stimulation with a modulation of the firing rate that spontaneously returned to baseline levels within 121 ms. This transient response indicates that electrical stimulation brings organoids in a state that cannot be sustained for a prolonged time and is, thus, unstable. This contrasts with previous findings in brain organoids^46^ as well as with studies on ES-induced state changes in brain slices^55^, but it aligns with an extensive literature of transient ES responses observed across species^9,56^.

Organoid units responded to ES mainly with a monophasic excitatory pattern, and, to a lesser amount, with a monophasic inhibitory or biphasic mixed pattern. This sharply contrasts with the tri-phasic patterns reported in-vivo in the premotor cortex of mammals during wakefulness ^9^. The monophasic excitatory response to ES in organoids closely resembles the one elicited in ferret brain slices in a slow oscillation regime and in cultured cortical networks^56,57^. Instead, the biphasic mixed pattern is reminiscent of the ES responses elicited in-vivo during sleep, anesthesia^58,59^ or in slices during pharmacological manipulations replicating awake activity^56^. The three phases of the prototypical response to ES encompasses an early excitatory phase, followed by an intermediate inhibition, and a late excitation.

The early excitatory phase, shown across all models including our organoids, is attributed to an activation either due to direct ES activation or polysynaptic excitation^60^. The observation of this early excitatory response in cortical organoids demonstrates that ES does not just modulate organoids maturation^61^, but induces also a transient modulation of their electrical activity above the spiking threshold, potentially recruiting polysynaptic, circuit-level dynamics. The intermediate spiking inhibition observed in several models may derive from different factors, including active inhibition mediated by inhibitory neurons^55^, refractoriness^62^, or the withdrawal of excitatory inputs from subcortical drivers^9^. In our model, we observed the intermediate spiking inhibition only in a limited subset of neurons. Thus, we can hypothesize that the mechanism mediating inhibition may not be fully developed in organoids and enables the early activation to linger for a longer time and last up to 400 ms. Our model changes two of the potential sources of intermediate inhibition compared to intact in-vivo brains. First, our organoids recapitulate a developmental stage in which GABA, which is the main inhibitory neurotransmitter in the adult cortex, has an excitatory effect on downstream neurons due to high intracellular chloride concentration^63,43^. Second, cortical organoids lack the thalamus, so that they do not experience the withdrawal of excitatory inputs induced by thalamic inhibition^9^. Thus, the reduction in the inhibitory phase observed in our dataset may depend on both the reduction of intracortical inhibition and the non-withdrawal of subcortical excitatory drivers.

Finally, the late excitatory phase has been ascribed to an excitatory feedback originating from the thalamus^9^. As such, the lack of a late excitatory phase in our cortical organoids provides additional, causal evidence supporting the thalamic origin of this late spiking response to ES.

Space-wise, we found that the magnitude of both excitatory and inhibitory responses to ES was negatively correlated with the distance from the ES site. This aligns with previous studies reporting analogous results^2^ and suggests that electrical stimulation is maximally effective on neurons in its immediate vicinity^16,64^. This is likely due to the high concentration of charges nearby the ES site, which is maximally effective on close, both excitatory and inhibitory, neurons. Assuming that organoid neurons connect following the distance based profile described in other models^65,66^, distant neurons receive less ES charges and are separated by more synapses from the stimulation site, leading to a negative relationship between response magnitude and distance from the ES site.

Latency-wise, we observe local spiking responses that onset 21 ms after the stimulus, a substantial delay compared to the 4 ms reported in mice^59^. Notably, the latency that we observe in organoids exceeds even the 11 ms latency of spikes mediated by transsynaptic activation in mice. Albeit some very early spiking responses occurred before 5 ms^67,68^ may have been masked by the artifact-removal procedures, the fact that spiking modulations emerge, on average, 21 ms after ES suggests that organoid neurons may need a longer time to initiate the stimulus-induced spiking response. This is in line with previous works showing that immature neurons, that represent at least part of the cells in our organoids, exhibit a delayed spike onset time^69^. Accordingly, organoid cells exhibit substantial variability in their developmental stage and, consequently, in their latency to generate spiking responses. This may explain the considerable variability in the response onset, ranging from 5 to 155 ms, and potentially, the emergence of intermediate excitatory responses.

In an effort to establish a reference to explore organoids responses to ES in future studies, we analyzed how they vary with the number of trials and across neurons. Using our default set of ES parameters, we found that 26 trials are sufficient to achieve a reliable response to ES, and that more trials may just marginally improve the quality of the average response in terms of cosine similarity. The average of 26 trials led to a response with an MSE of 3% compared to the response obtained using 120 trials, and the two were highly correlated over time. Depending on the specific goal of the data acquisition and the related downstream analyses, distinct criteria can be used, such as 5% of the MSE, achieved after 17 trials, or 95% of the Spearman correlation, achieved after 60 trials. Using this dataset, we also found that organoid neurons exhibit a substantial variability over time, and that ES quenches their variability in a rapidly evolving fashion. While spiking variability is reduced for 280 ms after ES, only the first 20 ms are highly stereotyped across neurons, pointing at prolonged and rapidly evolving network-level integration of information. This aligns with previous works showing that peripheral stimuli quench neural variability for prolonged time windows^54^. The different time courses of variability between our and previous experiments can be explained by their different models and stimulations used in the studies. On one hand, ES is an artificial stimulation that elicits highly non-physiological response patterns, compared to sensory stimulations. On the other hand, cortical tissue may need a certain level of maturation, beyond the one of the organoids included in our study, to develop its ability to integrate information. A further possibility is that inter-area interactions, that are absent in individual cortical organoids, may mediate stereotyped responses over time. Based on previous literature, we can speculate that ES in the intact mouse brain elicits stereotypical responses across all neurons nearby the stimulation site with several peaks over 200 ms^9,59^. Conversely, the immature cortical microcircuits in organoids are sufficient to sustain long-lasting stereotypical responses to ES, but they are not sufficient to induce late peaks of stereotypical activity, which may require further maturation or inter-area connections.

Our work provides a systematic study of how distinct ES parameters influence the spiking response to ES in cortical organoids. We found that the response spiking rate is negatively correlated with pulse intensity and duration but not correlated with stimulation frequency and charge density. Interestingly, this finding contrasts with recent studies in organoids showing a positive correlation linking ES intensity and duration with spiking responses in connected organoids stimulated with monophasic pulses^70^ or in individual organoids after patterned voltage stimulation^46^. Several factors may explain this difference, such as the use of distinct protocols for generating and connecting organoids, the exploration of ES protocols using monophasic or voltage pulses, the different focality of the stimulating pulses, and the use of different stimulating and recording systems. Our findings, compared in the same way to several studies in humans, in which higher pulse intensity and duration were associated with larger high frequency activity (HFA) responses^13–16^, a proxy to infer spiking activity in LFP recordings^71^.

While this unique model provides key insights on the effects of ES on cortical circuits, several limitations need to be considered in the interpretation of these results. ES can induce electrical artifacts in MEA recordings^72^. We developed and applied dedicated procedures to minimize the impact of these artifacts (see methods), verified that the signal was not saturated around the stimulus, and ensured that the spikes observed before and after the stimulus were unchanged and reflected biological spike waveforms. Nevertheless, we cannot fully exclude that these artifacts may have masked spikes occurring early after ES, and thus that ES may drive even larger or earlier spiking responses.

Furthermore, our study was performed on organoids in which potassium channels were blocked using a 4AP solution. Future studies are needed to reveal how these results translate to brain organoids without 4AP or with other pharmacological manipulations.

Our experiments were performed over a time-window of 40.26 ± 3.01 minutes. While this duration may not be enough to capture potential fluctuations in neuronal excitability over hours^73^, our experimental design accounts for potential fluctuations of excitability over minutes by comparing responses obtained within the same parameter block. Further experiments are needed to explore whether the excitability of brain organoids fluctuates over longer time scales.

For each parameter, our work explored a set of values that aligns with previous literature but does not exhaust the broad range of values used in previous studies. This is especially true for stimulation frequency, which we explore only up to 3 Hz. While we were limited to this maximum frequency due to technical limitations of our hardware, future work is needed to explore the effects of higher stimulation frequencies, as this can provide key insights to guide their increasing use of high frequency stimulation in clinical settings^74^.

With the exception of the default parameter set, our experiments were limited to 30 stimuli per each parameter set per organoid. While these values align with previous studies investigating the effects of ES on human brain activity^15,16^and our trial-based analysis confirms that this number of trials is sufficient to achieve reliable ES responses, more stimuli may have allowed us to identify neurons responding with more subtle firing rate modulations. Future studies can build on these findings to identify whether and to what extent higher trial numbers will capture more ES responses.

## Notes

**Conflicts of interest:** D.N. and N.S. are junior scientists of Manava Plus, F.F. and R.F. are senior scientists of Manava Plus, S.R. is the Chief Medical Officer of Manava Plus.

### Competing Interest Statement

D.N. and N.S. are junior scientists of Manava Plus, F.F. and R.F. are senior scientists of Manava Plus, S.R. is the Chief Medical Officer of Manava Plus.

